# Engineered spermidine-secreting Saccharomyces boulardii enhances olfactory memory in Drosophila melanogaster

**DOI:** 10.1101/2025.01.30.635726

**Authors:** Florance Parweez, Roger Palou, Ruizhen Li, Lanna Kadhim, Heath MacMillan, Mike Tyers, X. Johné Liu

## Abstract

The polyamines putrescine, spermidine and spermine are ubiquitous metabolites synthesized in all cells. The intracellular levels of polyamines, especially spermidine, decrease in aging. Oral spermidine supplementation has been reported to alleviate aspects of age-related disease in animal models, including decline in learning and memory. The diverse health benefits of spermidine supplementation, often at doses that do not significantly alter spermidine levels of target organs, suggests that exogenous spermidine may have a common site of action, the gastrointestinal (GI) tract. To directly deliver spermidine to the GI tract with minimum impact on the global spermidine levels, we engineered the probiotic yeast *Sacchromyces boulardii* (Sb) to overproduce and secrete spermidine. We tested the effects of a spermidine-producing yeast strain (Sb576) on age-associated learning and memory decline in an olfactory classical conditioning in *Drosophila melanogaster*. Feeding of newly emerged adult flies [w^1118^(isoCJ1)] for 30 days with food supplemented with live Sb576, but not live wild-type Sb (SbWT) or free spermidine, reduced age-associated short-term memory (STM) decline. Notably, Sb576 supplementation, but not SbWT or spermidine supplementation, of either young flies or old flies for only three days also enhanced STM without affecting locomotive ability. Transcriptome analysis of the gut revealed relatively few (30) differentially overexpressed genes in the Sb576 group compared to the SbWT group including the gene coding neuropeptide Dh31, which has been implicated in memory in the flies. These results demonstrate that in situ production of spermidine by a synthetic biotic yeast in the GI tract can enhance STM, and further suggest a mechanism involving the gut-brain axis.

## Introduction

The polyamines putrescine, spermidine and spermine, are produced in all organisms from bacteria to humans. In eukaryotic cells, ornithine decarboxylase (ODC) converts L-ornithine to the diamine putrescine. To synthesize the triamine spermidine, two additional enzymes are required. S-adenosylmethionine (SAM) decarboxylase (SAMDC) converts SAM to decarboxylated SAM (dcSAM). Spermidine synthase transfers an aminopropyl group from dcSAM to putrescine to form spermidine. Similarly, spermine synthase transfers an aminopropyl group from dcSAM to spermidine to form the tetraamine spermine ^1^. Polyamines are essential metabolites since deletion of the *Odc* gene causes peri-implantation embryonic lethality in mice^2^. It has been reported that the level of polyamines, especially spermidine, decreases in many mammalian tissues during aging ^3^. Recent attention has been focused on the potential of spermidine supplementation as an anti-aging strategy, through extension of lifespan in animal models and amelioration of inflammatory bowel disease, colon cancer, cardiovascular disease, and neurodegeneration ^4,5^. Specifically, free spermidine supplementation in *Drosophila* food extends lifespan ^4^ and prevents aging-related olfactory learning and memory decline ^6^.

Various molecular mechanisms have been proposed to explain the health benefits of spermidine supplementation. Biochemically, spermidine serves as a precursor for hypusine, a functionally necessary lysine adduct only found in eIF5A ^7^, a translation elongation factor essential for overcoming ribosome stalling at polyproline sequence stretches ^1^. Hypusination of eIF5A has been proposed to be responsible for spermidine effects on B cell rejuvenation in aging ^8^, reducing gut inflammation ^9^, fasting-mediated autophagy and longevity ^10^ and reducing brain aging in Drosophila ^11^. It has also been proposed that spermidine activates mitochondrial trifunctional enzyme in T cells to boost anti-tumor immunity ^12^ and/or inhibits T cell tyrosine phosphatase to reduce gut inflammation ^13,14^.

The reported diverse health benefits of spermidine supplementation are often at doses that do not significantly change intracellular spermidine levels in target organs, suggesting that spermidine might have a common site of action, namely the gastrointestinal (GI) tract. The GI tract houses the body’s largest immune ^15^ and endocrine ^16^ systems. Spermidine may exert systemic effects by protecting the intestinal epithelium ^17^ and modulating the immune and endocrine systems. On the other hand, long-term and systemic spermidine supplementation may carry health risks since high levels of spermidine has been associated with cancers and stroke ^18,19,20^. To directly deliver spermidine efficiently to the GI tract with minimum effects on systemic spermidine levels, we have engineered the probiotic yeast *Saccharomyces boulardii* (Sb) to overproduce and secrete spermidine. This Sb strain, called Sb576, harbors a deletion of the *OAZ1* gene, which encodes the antizyme inhibitor of Spe1 (Odc1 in yeast) and overexpresses *SPE1*, *SPE2* (which encodes yeast Samdc) and TPO5 (which encodes a yeast polyamine exporter). Sb576 retains viability in the gut lumen when administered orally to mice, increases GI lumen spermidine levels, and ameliorates ulcerative colitis and colon cancer in mice (Mohaqiq et al. DOI: <u>10.1101/2025.01.19.633601</u>). In this study, we tested the impact of food supplementation of Sb576 on olfactory learning and memory in *Drosophila*. We found that supplementation of Sb576, but not wild-type Sb (SbWT) or free spermidine (1mM) reduced aging-related short-term memory (STM) decline. Moreover, we found that short-term (3 days) supplementation of Sb576, but not SbWT or spermidine, enhanced STM in both young flies and aged flies.

## Results

### Long-term supplementation of Sb576 reduces aging-associated short-term memory decline

To explore the efficacy of the spermidine secreting strain Sb576 in delaying aging-related memory decline, we employed the *Drosophila* olfactory learning and memory model ^6^. Synchronized adult flies [w^1118^(isoCJ1)] ^21^ were fed on fly food (cornmeal, CM) supplemented with live SbWT, Sb576, or with 1mM spermidine (see Methods for details) for 30 days. Thirty-day-old flies were subjected to an olfactory learning and memory (STM) test according to the protocol of Malik and Hodge ^22^, using young flies (3-day-old) as a control. As expected, we found that 30-day-old flies fed on CM food had significantly lower performance index (PI) compared to young flies (Fig. 1A, young vs CM). Supplementation of CM with SbWT or with spermidine (SPD, 1mM) did not impact PI. However, Sb576 supplementation significantly improved PI, not only compared to CM-fed flies, but also compared to SbWT-supplemented flies and, to a slightly lesser degree (p=0.05, one-way ANOVA and Tukey test; p=0.01 by t test), to spermidine-supplemented flies (Fig. 1A). We also subjected the flies to locomotive ability testing and found that in general young flies performed better than 30-day-old flies. However, there was no significant difference in locomotive ability amongst the 30-day-old flies regardless of the food supplementation regime (Fig. 1B). We extracted polyamines from whole flies and performed LC-MS/MS analyses and quantification (Fig. 1C). These data revealed significant differences in spermidine levels amongst the groups (p=0.027, one-way ANOVA). Specifically, 30-day old flies (CM) had reduced spermidine levels compared to young flies. Spermidine supplementation, and Sb576 supplementation to a lesser degree, restored spermidine levels. In contrast, there was no significant difference in spermine (SPM) levels in any of the groups. Supplementation with Sb576 and with spermidine also significantly increased putrescine (PUT) levels compared to SbWT-supplemented group, although the overall putrescine levels were low and there was no difference between young and old flies (young vs CM). The increase in putrescine levels in flies supplemented with spermidine had been observed previously ^4^, indicating that catabolic conversion of spermidine to putrescine ^23^ is active in flies. Together, these results suggest that Sb576 supplementation reduces aging-associated STM decline without affecting aging-associated locomotive decline in *Drosophila*. We note that despite restoring spermidine levels in 30-day old flies to those found in young flies (Fig. 1C), free spermidine supplementation failed to reduce aging-associated STM decline (Fig. 1A), in contrast to a previous report ^6^.

**Figure 1.**
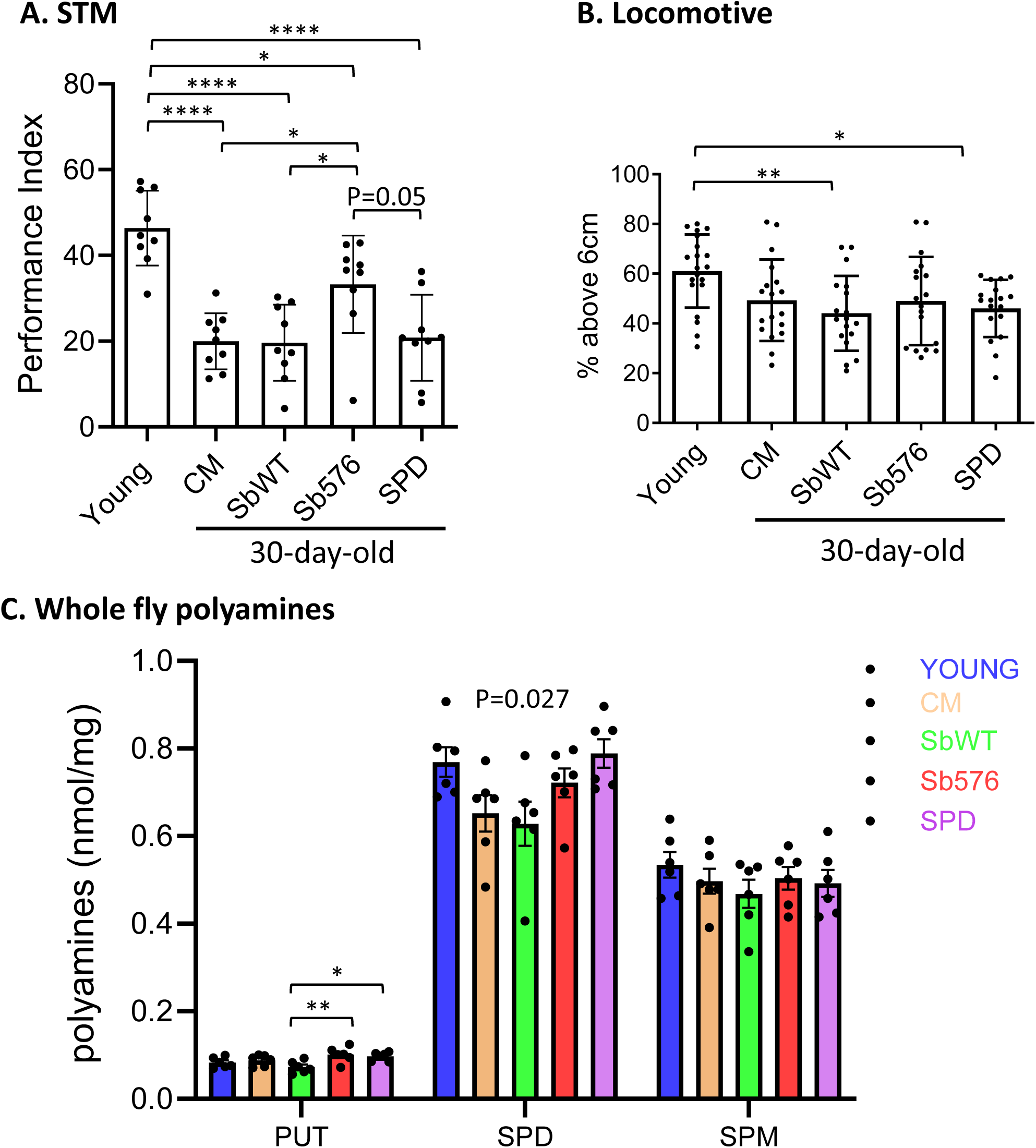
Sb576 reduces aging-associated STM decline in flies **A.** STM performance scores of young flies and flies after 30-day feeding on CM without supplementation or supplemented with live SbWT or Sb576. **B.** Locomotive ability test of the same flies immediately following the olfactory memory test in A. **C.** Whole body polyamine levels in young flies and 30-day old flies fed on CM or CM plus the indicated supplementation. Shown are means with SEM. P value represents One-way ANOVA and Tukey’s multiple comparisons test.

### Short-term Sb576 supplementation enhances STM in young flies and old flies

Given the specific effects of Sb576 on aging-related STM decline, but not on aging-related locomotive decline, we wonder if the Sb576 strain might impact STM independently of aging processes. To test this, we compared STM performance of newly emerged adult flies fed for three days with various live yeast supplements. For these experiments, we used CM food without baker’s yeast (CM^-Sc^) and supplemented with live baker’s yeast (*Saccharomyces cerevisiae*, Sc, strain Sigma 1278b), SbWT or Sb576. Newly emerged adult flies were transferred to CM^-Sc^ with the indicated live yeast supplementation and cultured for three days followed by STM test. We found that Sb576 supplementation significantly improved STM performance compared to supplementation with either ScWT or SbWT (Fig. 2A). No difference was found in any group in a locomotion test (Fig. 2B). To rule out the possibility that Sb576 may alter odor perception or response to electric shock used in conditioning and testing for STM assays, we performed odor acuity and shock reactivity tests and found no difference among the three groups of flies (Table 1). In separate experiments, we also compared young flies fed for three days on CM or CM containing 1mM spermidine and found no significant difference in STM performance (Fig. 2C) or locomotive ability (Fig. 2D) between the two, confirming results by others ^6^. Taken together, our results suggest that Sb576 enhanced STM performance but not locomotive activity in young flies.

**Figure 2.**
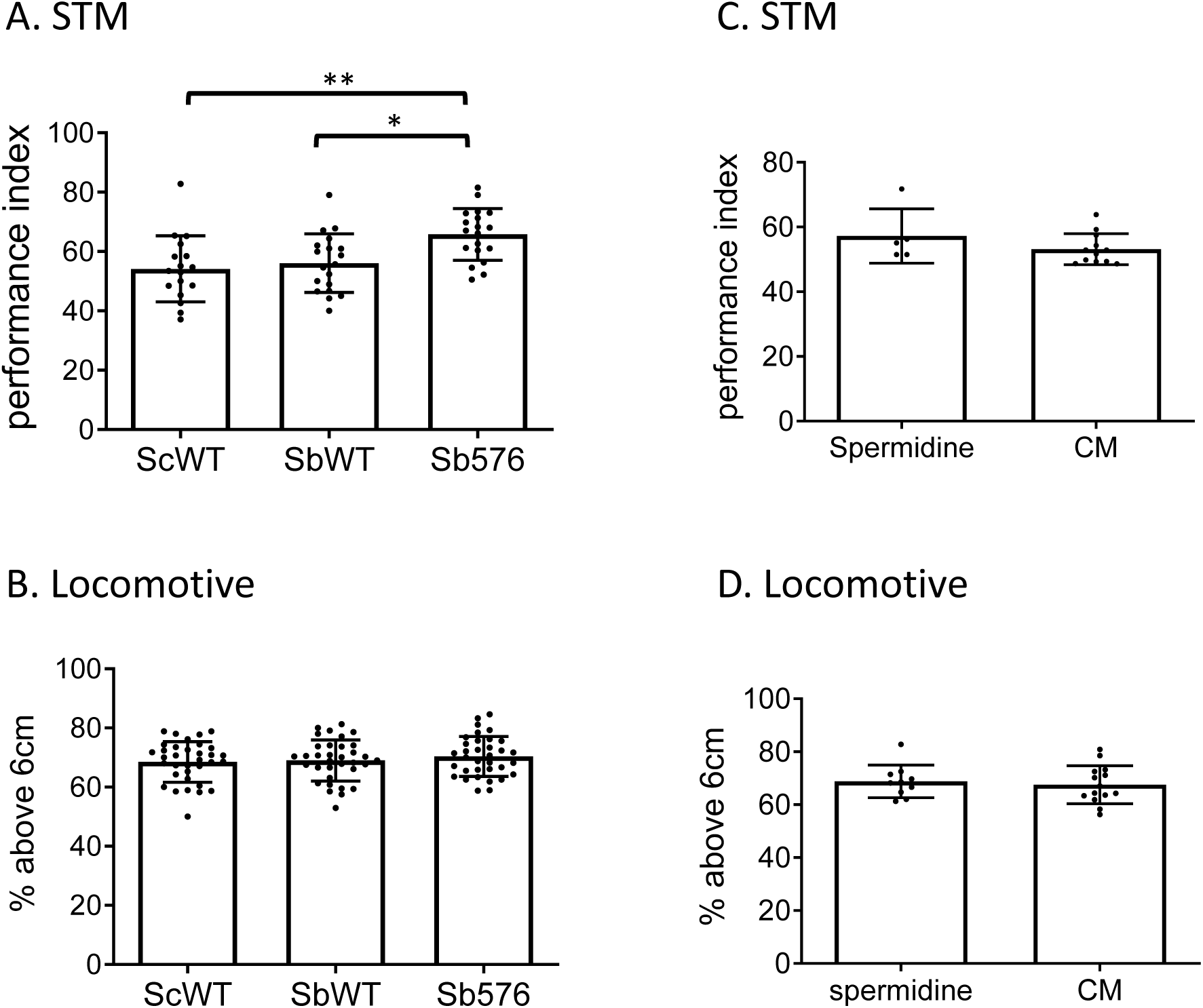
Sb576 enhances STM in young flies **A,B.** Newly emerged adult flies after three-day feeding on food supplemented with live ScWT, SbWT or Sb576 were tested for olfactory memory (A) and locomotive ability (B). Shown are means with SEM. Significant difference in memory scores was found between Sb576 and ScWT, and between Sb576 and SbWT (One-way ANOVA and Tukey’s multiple comparisons test). **C,D.** Newly emerged adult flies after three-day feeding on cornmeal food or cornmeal food plus 1mM spermidine were tested for olfactory memory (C) and locomotive ability (D). Shown are means with SEM.

**Table 1.** Odor acuity and shock reactivity Performance index (means with SEM, n=3) of flies fed on food supplemented with the indicated live yeast. Newly emerged adult flies were fed for three days before testing for odor acuity or four days before testing for electric shock reactivity.

|  | Olfactory Avoidance |  | Shock Reactivity |
| --- | --- | --- | --- |
|  | MCH | OCT |  |
| ScWT | 62.9 ± 2.5 | 83.4 ± 8.2 | 82.6 ± 4.4 |
| SbWT | 66.8 ± 8.4 | 84.6 ± 3.5 | 71.9 ± 6.8 |
| Sb576 | 67.5 ± 9.8 | 80.0 ± 7.7 | 67.7 ± 13.4 |

We reasoned that if Sb576 can enhance STM in young flies, independently of aging process, it may also enhance STM performance in aged flies that exhibit an age-associated memory deficit^24^. We cultured flies on regular CM food for 27 days followed by feeding for three days on CM alone, or CM supplemented with spermidine (1mM), or CM^-Sc^ supplemented with live SbWT or Sb576. All flies were subjected to the same STM test as 30-day-old flies as above, with young flies as a control. Twenty-seven-day-old flies treated with Sb576 supplementation for three additional days performed significantly better than the other 30-day-old control groups fed on regular CM or supplemented with SbWT or spermidine. Despite the improvement in STM of old flies fed Sb576, we observed a significant difference in STM performance between the Sb576 group and young flies (Fig. 3A), indicating that the 3-day treatment with Sb576 only partially rescued the memory deficit lost during aging. In contrast to STM performance, Sb576 supplementation did not alter locomotive activity in old flies (Fig. 3B).

**Figure 3.**
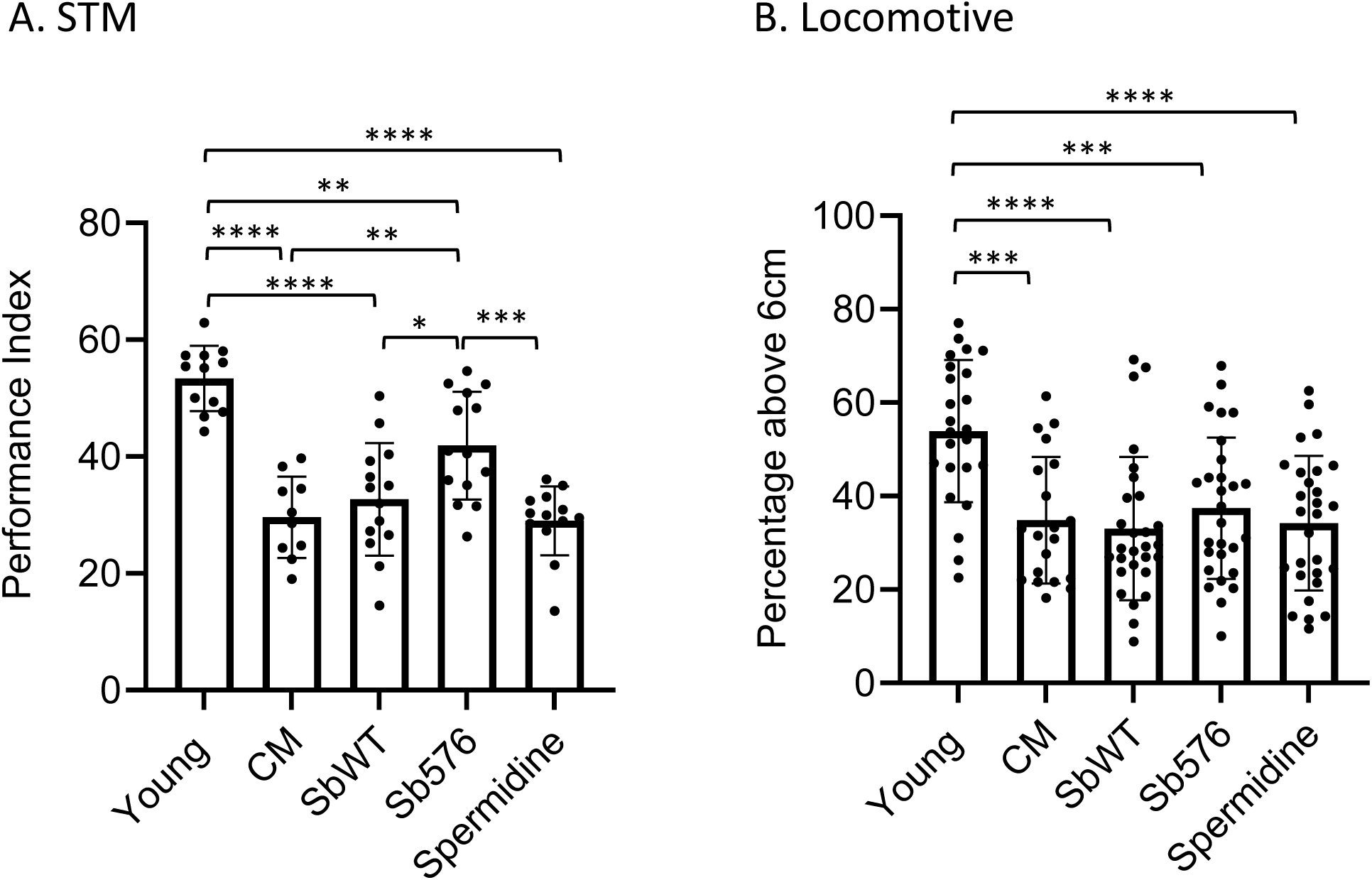
Sb576 enhances STM in aged flies Newly emerged adult flies were fed on cornmeal food for 27 days before transferring to yeast-free cornmeal food supplemented with live SbWT or Sb576, or to cornmeal food plus 1mM spermidine. Flies were tested three days later for olfactory memory **(A)** and locomotive ability **(B)**. Shown are means with SEM. Significant difference in olfactory memory was found between flies fed on Sb576-supplemented food and those on CM or SbWT-or spermidine-supplemented food.

### Intestinal transcriptome analysis of flies treated with Sb576 vs SbWT

The impact of short-term 3-day Sb576 supplementation on STM suggested a potential direct mechanism of action involving the gut brain axis. To explore this possibility, we carried out RNAseq analysis of intestines from young versus SbWT-or Sb576-treated old flies (i.e., those shown in Figure 1). Whereas numerous differentially expressed genes were found between young flies and the two groups of old flies, few such differentially expressed genes were found between SbWT-and Sb576-treated old flies. Specifically, only 30 genes with GFOLD values ^25^ greater than 1.5 were found over-expressed in Sb576-treated group compared to SbWT-treated group. All three yolk protein genes, Yp1-3, were among this group, suggesting a positive impact of Sb576 administration on egg production and fertility ^26^. Most interestingly, the diuretic hormone Dh31 was highly overexpressed in Sb576-treated old flies compared to SbWT-treated old flies (GFOLD=1.74), and in fact, Dh31 was overexpressed in Sb576-treated old flies compared to young flies (Table 2). Dh31 is one of 10 neuropeptides known to be expressed in enteroendocrine cells in the *Drosophila* intestine ^27–29^, and was the only such gene over-expressed in Sb576-treated flies (Table 2).

**Table 2.** Expression of midgut hormones Relative (mRNA) expression levels of neuropeptide genes known to be expressed in *Drosophila* enteroendocrine (EE) cells in the intestinal tissues of young flies, SbWT-or Sb576-treated old flies.

| EE-expressed<br>Neuropeptide genes | RPKM<br>Old-SbWT | RPKM<br>Old-Sb576 | g-fold<br>(Sb576/SbWT) | RPKM<br>Young |
| --- | --- | --- | --- | --- |
| Dh31 | 4 | 21 | 1.74 | 7.9 |
| Akh | 0.2 | 1.9 | 1.05 | 15 |
| TK | 42 | 52 | 0.05 | 40 |
| NPF | 27 | 26 | -0.01 | 38 |
| AstA | 37 | 41 | 0 | 7.8 |
| AstC | 103 | 104 | -0.10 | 58 |
| Orcokinin | 26 | 29 | 0 | 25 |
| MIP | 0.8 | 0.9 | 0 | 0.6 |
| CCHa1 | 41 | 33 | 0 | 7.4 |
| CCHa2 | 38 | 45 | 0 | 26 |

## Discussion

We have demonstrated that supplementation of *Drosophila* food with an engineered, spermidine-secreting Sb strain, Sb576, reduced aging-related decline in learning and memory (STM). Moreover, we have shown that short-term (3 days) Sb576 supplementation enhanced STM in young flies, as well as in aged flies with memory deficit. The memory enhancement was unique to Sb576, and not shared by either SbWT or free spermidine (1mM) supplementation. The inability of spermidine supplementation to reduce aging-associated memory decline contrasted with previous finding that spermidine supplementation (1mM or 5mM) reduced aging-associated decline of both STM and intermediate term memory (ITM) ^6^. Although the reason for this apparent discrepancy is not known, we note that in previous studies spermidine supplementation was initiated during parental fly mating and then continued through egg-to-adult development and adult aging ^6,11,30^. It is therefore possible that the beneficial effects of free spermidine supplementation reported previously require prolonged administration over the entire lifespan of the animal, starting from fertilized eggs ^6^. We note that our spermidine supplementation regime was sufficient to restore spermidine levels in old flies to the level found in young flies, as efficiently, if not more so, as Sb576. Thus, the inability of direct spermidine supplementation to reduce memory decline in aged flies in our study was unlikely to be due to insufficient spermidine supplementation.

Polyamines, especially spermidine and spermine, are potent enhancers of N-methyl-D-aspartate (NMDA) receptor signaling through potentiation of ligand binding to the receptor ^31^. However, the effect of exogenous spermidine supplementation on learning and memory may be complex. Long-term oral spermidine supplementation improves brain functions in flies ^6,11,30^ and mice ^32,33^. Since spermidine has very limited permeability through the blood-brain barrier ^34,35^, the effects of spermidine on brain functions in these long-term supplementation experiments may be indirect. Consistent with this interpretation, spermidine levels have been found to decline in some mouse tissues but not in the brain during aging ^3^. Indeed, Rushaidhi et al., ^36^ found that spermidine levels increase in some hippocampal subregions and do not change in others during rat aging ^36^. Short-term intrahippocampal infusion of spermidine after training improves brain functions in rats, likely through direct modulation of NMDA receptors in postsynaptic neurons ^37–39^. Intracerebroventricular injection of drugs that inhibit spermidine synthesis or disrupt spermidine-NMDA receptor binding counteracts β Amyloid peptide-induced memory impairment in mice, likely through modulating extrasynaptic NMDA receptors ^40^.

Since spermidine supplementation, which was as effective, if not more so, as Sb576 supplementation in restoring spermidine levels in aged flies, did not impact STM in flies, it seems unlikely that the effects of Sb576 on STM occur through direct interaction of spermidine with brain NMDA receptors in the flies. Furthermore, Sb576 supplementation not only reduced aging-related memory decline, but short-term (3 days) Sb576 supplementation also enhanced memory of young flies, as well as of aged flies that already suffered significant memory deficits. We propose that Sb576 enhances memory in the flies through a mechanism that relies on the gut-brain axis. In support of this hypothesis, our transcriptome analysis of *Drosophila* intestine revealed that diuretic hormone, Dh31, was one of the most highly overexpressed genes specific to Sb576 administration. Dh31 is expressed in neurons of the brain and ventral nerve cord and in enteroendocrine cells of the midgut, and notably midgut expressed Dh31 controls nutrient-mediated behavior ^27^. Interestingly, a recent study ^41^ demonstrated that neuron derived Dh31 is required for ITM but not STM, raising the possibility that midgut derived Dh31 may be involved in Sb576 enhancement of STM, a possibility requiring future investigation. Collectively, our studies with spermidine secreting Sb in flies position this synthetic biotic platform for evaluation of effects on learning and memory in more complex model systems.

## Methods

### *Drosophila* rearing and food preparation

Flies were reared in standard *Drosophila* cornmeal food (CM; recipe for one liter in water: agar, 6.7g; dry yeast baker’s yeast, 23.1g; NaKT or potassium sodium tartrate tetrahydrate, 6.30g; CaCl_2_, 0.53g; sucrose, 22.8g; dextrose, 45.5g; cornmeal, 55.1g; 8.5mL of acid mix containing 41.8% proprionic acid and 4.2% of phosphoric acid). Food components and water, minus the acid mixture, were added to a rice cooker, and the mixture was brought to boiling with stirring to dissolve components and then kept at boiling for 20 minutes. After the stock had cooled down to 60-65^0^C, the proprionic acid and phosphoric acid mixture was added and mixed thoroughly. For spermidine supplementation, spermidine trihydrochloride (Sigma Aldrich; made as a 1M stock in water and stored at -20^0^C in single-use aliquots) was added to CM to a final concentration of 1mM and mixed thoroughly before pouring into bottles. For live yeast supplementation, we prepared CM for 30-day supplementation experiments or CM without dry yeast (CM^-Sc^) for 3-day experiments, as indicated in figure legends. The parental *S. boulardii* strain MYA-796 was obtained from ATTC. The spermidine secreting strain Sb576 was built in MYA-796 by deletion of *OAZ1* and integration of a multi-genic expression cassette at a *TY2* locus, with the following genotype: *oaz1Δ TY2::pTDH3-TPO5-FLAG pTDH3-SPE1-2A-SPE2-FLAG-KanMX::Ty2*. The control *S. cerevisiae* strain Sigma 1278b was as described previously ^42^. Live yeast were grown to saturation in rich YPD medium, added to a final density of 5×10^7^ cells per mL CM or CM^-Sc^ (at 50-55^0^C), and mixed thoroughly before pouring into bottles. All flies were stored in an incubator at 25^0^C, with humidity at ∼50% and a 12h:12h light: dark cycle.

### Olfactory learning and short-term memory (STM) assays

STM assays were carried out in a small room set at 25^0^C, 60% relative humidity, and under dim red light. Flies were placed into the room 24h prior to the test to acclimatize. The Fly Training Machine System, or T-maze was purchased from CelExplorer (https://www.celexplorer.com/product_list.asp). Flies (50-100 flies per group) were placed into the training tube and exposed to fresh air (at 2.0L per min flow rate) for 90s, followed by exposure to one odor (conditioned stimulus, CS^+^, 0.15% 3-octanol [Sigma Aldrich] or 0.12% 4-methylcyclohexanol [Sigma Aldrich]) simultaneously with electric shock (60V/1.5s with 3s intervals for a total of 60s). Flies were then given 30 seconds to rest before being exposed to the second odor (CS^-^, 4-methylcyclohexanol or 3-octanol) without shock for 60s. During the testing period, flies were moved to the choice chamber where they were exposed simultaneously to the two odors (CS^+^ and CS^-^), on each side of the chamber, and allowed 120s to choose a side. Flies were trapped in their respective choice tubes and counted. The performance index was calculated as the number of flies avoiding the shocked odor (CS^+^) minus the number of flies avoiding the non-shocked odor (CS^-^) divided by the total number of flies in the test and multiplied by 100 ^22^. For each PI value, two groups of flies were tested, one using 3-octanol as CS^+^ and the other using 4-methylcyclohexinol as CS^+^, and the average of the two sub-PIs was taken as PI (n=1).

### Locomotive activity test (climbing Assay)

The same flies used for STM assays were used subsequently for the climbing assay on the same day. The two groups of flies for n=1 in the STM test were tested separately in climbing assays, resulting in twice the sample size (n). Flies were transferred into a *Drosophila* culture vial with a line marked 6cm from the bottom, gently tapped the flies down to the bottom, and video-captured while they climbed up the side of the tube. Video was paused at the 10s-time frame. Flies below and above the 6cm line were counted from the captured image at the 10s-time frame and presented as % above the 6cm line.

### Olfactory acuity

Odor avoidance responses to 3-octanol and 4-methylcyclohexanol was measured by giving a group of untrained flies the option to choose between 1.5% 3-octanol, or 2% 4-methylcyclohexanol, vs fresh air (all at 2.0L/min). After 120 seconds, flies were trapped in their respective choice tubes and counted. Performance index was calculated as the number of flies collected in fresh air arm minus that in the odor arm, divided by total number and multiplied by 100 ^22^.

### Shock reactivity

Shock reactivity was quantified by placing a group of untrained flies into the T-maze where one side of the maze had a training tube where the shock (60V/1.5s with 3.5s intervals for a total of 60s) was delivered. The other side of the maze had the tube but not shock. After 1 minute, flies were trapped into their respective tubes and counted. The performance index was calculated as number of flies recovered from the non-shock tube minus that from the shock tube, divided by the total number and multiplied by 100 ^22^.

### Polyamine determination

Polyamines were extracted from whole flies according to Qin, et al. ^43^ with slight modifications. Briefly, 30 adult flies (15 male and 15 female) were frozen in liquid N_2_ and ground using a disposable pestle. Boiling water (495 μL) was mixed with 5 μL of 100μg/mL deuterated spermidine, Spermidine-(butyl-d_8_) trihydrochloride (709891; Sigma-Aldrich), and the mixture was added to each ground sample. The tube was then vortexed vigorously to suspend the ground sample and kept boiling for 30 min. Tubes were then placed on ice for 5 minutes. The post centrifugation (twice at 15,000 rpm at 4^0^C,15 min) supernatant was then stored at –80^0^C or directly subjected to the quantification step.

All extracted samples were analyzed by using a validated liquid chromatography mass spectrometry/mass spectrometry (LC-MS/MS) method. The chromatographic separation was achieved on the Thermo Accela LC system with a Supelcosil AZB+Plus 3μ (100×2.1mm). Gradient elution (0.1% heptafluorobutyric acid as the ion-pairing agent with μl/ml at 40^0^C) was used to separate all polyamines and internal standard (Spermidine-(butyl-d_8_) trihydrochloride). Mass spectral analyses were accomplished on a quadrupole tandem Mass Spectrometry (TSQ Quantum Access MAX, Thermo). Nitrogen was used as curtain, nebulizer and collision gases. User controlled voltages, gas pressures, and source temperature was all optimized for the detection of polyamines and IS. MS was quantified using electrospray multiple reaction monitoring (MRM) in positive mode and the MRM transitions were m/z 89.1to 72.1 for putrescine, m/z 146.2 to 112.2 for spermidine m/z 203.2 to 129.1 for spermine, and m/z 154.1 to 120.3 for IS. 10 μL was used for injection and the autosampler tray temperature was 22^0^C. The effective linear range was 0.2 -5μg/mL for putrescine, 01-2.5µg/ml for spermidine, and 0,05-1 μg/mL for spermine. Inter-batch precision (CV%) varied between 6 and 18% and intra-batch accuracy varied between 87 and 118%. Validation results displayed that all polyamines and IS were stable for 10 hours 22^0^C.

### RNAseq

To eliminate/reduce yeast (and yeast RNA) in the intestine, flies were transferred to bottles containing filter paper saturated with 5% glucose in water for four hours before being killed by CO_2_ suffocation. The entire intestine excluding the crop was excised in PBS. Isolated guts were then placed into Eppendorf tubes. Total RNA was isolated from 30 female guts using Qiagen’s RNeasy Plus kit. RNA sample quality assessment was performed with the Fragment Analyzer Standard Sensitivity RNA assay (Agilent) and concentration measured with the Qubit 3.0 HS RNA assay (Thermo). NGS libraries were prepared with the Stranded mRNA library prep kit (Illumina) using 400 ng of total RNA input. Sequencing was performed with the Nextseq 2000 P1 100 cycle flow cell (Illumina). FASTQ sequencing file reads were aligned to the *D. melanogaster* genome BDGP6.46 genome, Ensembl transcriptome 59 using the nf-core/rnaseq pipeline ^44^ version 3.14.0. Fold change estimates between samples were generated using GFOLD ^25^ to compare STAR/salmon pseudocount values.

### Statistical Analysis

Statistical analysis and graphing were performed using GraphPad Prism software. All quantitative values are shown as means ± standard error of the mean (SEM). To compare group means, one-way ANOVA analysis was used followed by Tukey’s multiple comparisons test. *, p<0.05; **, p<0.01; ***, p<0.001; ****, p<0.0001.

## Acknowledgements

We thank Marshall W Ritchie (MacMillan lab) for advice on fly rearing and Guijun Zhang (OHRI) for polyamine determination. We thank Drs. Joshua Dubnau (Stony Brook University) for providing stocks of w^1118^(isoCJ1) flies. The authors would like to acknowledge the assistance of StemCore Laboratories Genomics Core Facility (OHRI, uOttawa), RRID:SCR_012601, and the Ottawa Bioinformatics Core Facility (uOttawa/OHRI), RRID:SCR_022466. This work was supported by grants from the Canadian Institutes of Health Research (CIHR) to XJL (PJT 168891) and MT (FDN-167277), and an ELEVATE award from Ottawa Hospital Research Institute to XJL.

## Author contributions

FP: Design and perform experiments, analysis of data and drafting manuscript.

RP: Design and perform experiments, analysis of data.

RL: Design and perform experiments, analysis of data.

LK: Perform experiments.

HM: Supervision of study.

MT: Conception and supervision of study, acquisition of funding and drafting the manuscript.

XJL: Conception and supervision of study, acquisition of funding and drafting the manuscript.

## Data availability statement

All data associated with this study are available in the main manuscript.

## Notes

### Competing Interest Statement

The authors have declared no competing interest.

